# The energetic cost of human walking as a function of uneven terrain amplitude

**DOI:** 10.1101/2024.11.08.622353

**Authors:** Seyed-Saleh Hosseini-Yazdi, Arthur D. Kuo

## Abstract

Humans expend more energy walking on uneven terrain, but the exact cost varies across terrains. Few experimental characterizations exist, each describing terrain qualitatively without any relation to others or flat ground. This precludes mechanistic explanation of the energy costs. Here we show that energy cost varies smoothly and approximately quadratically as a function of terrain amplitude. We tested this with healthy adults (N=10) walking on synthetic uneven terrain with random step heights of parametrically controlled maximum amplitude (four conditions 0 – 0.045 m), and at four walking speeds (0.8 – 1.4 m · s^−1^). Both net metabolic rate and the rate of positive work increased approximately with amplitude squared and speed cubed (*R*^2^ = 0.74, 0.82 respectively), as predicted by a simple walking model. The model requires work to redirect the body center of mass velocity between successive arcs described by pendulum-like legs, at proportional metabolic cost. Humans performed most of the greater work with terrain amplitude early in the single stance phase, and with faster walking late in stance during push-off. Work and energy rates changed with approximately linear proportionality, with a ratio or delta efficiency of 49.5% (*R*^2^ = 0.68). The efficiency was high enough to suggest substantial work performed passively by elastic tendon and not only by active muscle. Simple kinematic measures such as mid-swing foot clearance also increased with terrain amplitude (*R*^2^ = 0.65), possibly costing energy as well. Nevertheless, most of the metabolic cost of walking faster or on more uneven terrain can be explained mechanistically by the work performed.

**Summary statement:** Humans perform more work and expend more energy on uneven terrain, increasing with the square of terrain amplitude and the cube of walking speed.

## Introduction

Humans walk on uneven terrain with a variety of biomechanical adjustments that could potentially explain the greater energy they expend. The adjustments can be described by kinematic gait parameters such as step distances that change on uneven terrain, as do their respective variabilities [1], [2]. Such adjustments presumably stem from kinetic changes such as to the work and forces produced by muscles [1]. However, it is challenging to form a mechanistic connection between energy expenditure and other measures without knowing how each varies as a continuous function of the others. If uneven terrain could be varied continuously, it might enable better explanation of the greater energy expenditure.

Most kinematic gait measures are affected by uneven terrain. Humans take shorter, wider, and briefer steps on uneven terrain compared to same-speed flat terrain [1], [2]. Such changes to the overall gait pattern may account for some of the energy cost, as suggested by the fact that experimentally imposed changes to average gait parameters cause greater energy expenditure on flat ground [3], [4]. Humans also walk with greater step variability [1], [2], [5], which may reflect step-by-step adjustments such as for foot placement and lateral balance. Again, imposed perturbations to balance lead to greater energy cost during flat walking [6], [7]. Conversely, externally stabilizing lateral balance also leads to reduced step variabilities and energy cost [8], [9], [10]. Kinematic measures are helpful because they are relatively easy to measure, including in real-world settings [2], and can potentially correlate well with energy expenditure.

It is, however, preferable to develop a mechanistic basis for energy expenditure. One of the simplest mechanistic models predicts the work performed on the body center of mass (COM) [11], with an approximately proportional cost [12] that explains most of the energetics of level walking [13], [14]. This suggests a basic generalization of the same model to steps of uneven height, which induce fluctuations in timing and velocity that generally require more work to maintain the same speed [15], [16]. Of course, COM work is not the sole contribution, because the body also produces other work and forces peripheral to the COM or about the joints. Their associated costs are, however, thought to be smaller [11] and are less straightforward to predict. It is thus possible that COM work can account for much of the energetic cost of uneven walking, if it is a straightforward generalization of flat walking.

Both kinetics and kinematics are best characterized as a function of a controlled variable. Energy costs are greater on sand than brush [17] and on woodchips than sidewalk [2], but such observations are challenging to replicate or explain, because natural terrain varies in numerous unquantifiable and uncontrollable ways. One remedy is to experimentally control a single independent variable such as uneven terrain amplitude. Accordingly, we recently developed a method to vary terrain amplitude on a split-belt, instrumented treadmill [18], which facilitates kinetic and kinematic analysis. To ensure that properties such as ground stiffness and damping do not vary, the terrain may be synthesized from consistent material (polystyrene). This simplification also makes uneven walking amenable to modeling, which may suggest how terrain amplitude should theoretically affect the mechanics and energetics of walking.

The purpose of the present study was to test for three factors in uneven walking: kinetics, kinematics, and terrain amplitude. The principal hypothesis was based on kinetics, specifically that COM work would increase as predicted by mechanistic model, with a proportional contribution to energy cost. The second factor was kinematics, used to characterize how walking varies with terrain, perhaps in ways complementary to COM work. The third and final factor was terrain amplitude as a controlled variable potentially affecting kinetics, kinematics, and energetics. These three factors differ in how they are quantified, what they describe, and how they fit with a mechanistic model. They are therefore all potentially helpful for explaining the energetic cost of uneven walking.

## Materials and Methods

### Model predictions

We used a simple walking model to make rough predictions for work and energy cost (Figure 1). Briefly summarized, the model consists of pendulum-like legs with a point center of mass (COM) for the pelvis, and feet of infinitesimal mass [19], [20]. The COM moves in a series of pendular arcs, each redirected in a step-to-step transition by forces along the legs. The leading leg performs dissipative negative work in its collision (Figure 1B, CO) with ground. Active, positive work may be performed with an impulsive push-off (PO) by the trailing leg, supplemented if necessary by continuous hip extension during the single stance phase [15]. In steady, flat walking the push-off impulse ideally acts just prior to collision and with equal magnitude [19]. But on uneven terrain, the gait is generally not steady and the appropriate control strategy less obvious. An ideal would be to optimize the magnitude and timing of push-off over a fully anticipated sequence of uneven steps [21], but an uneven treadmill affords very limited ability to see and plan for upcoming steps. More feasible but less optimal strategies could include applying constant push-off or stance phase hip work each step, in amount sufficient to maintain overall speed [22].

**Figure 1:**
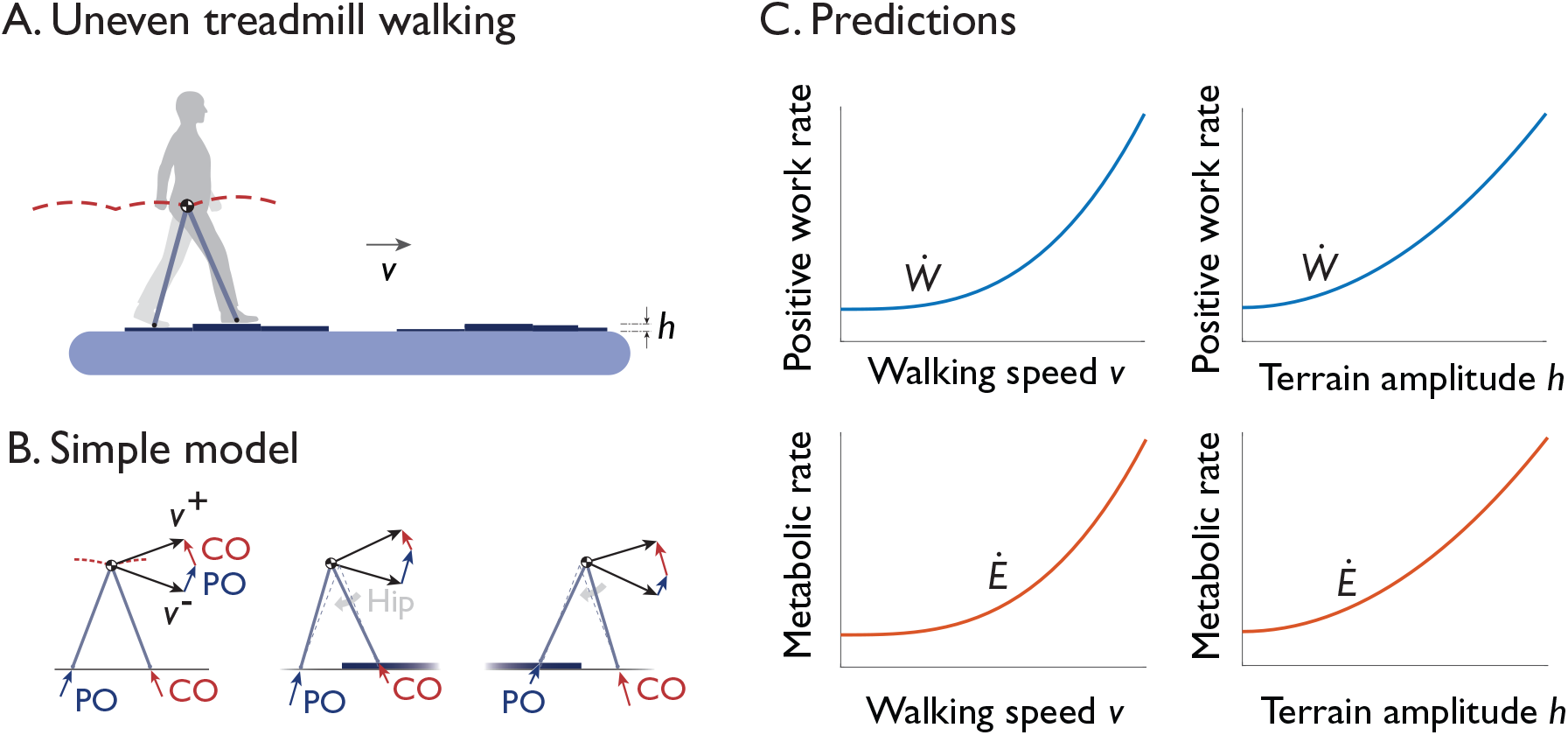
Uneven walking studied experimentally and theoretically. (A) In experiment, healthy adults walk on a split-belt, instrumented treadmill fitted with several different uneven terrain treadmill belts. The terrain is parametrically varied in terms of peak-to-peak amplitude *h*, and walking conditions by treadmill speed *v*. (B) A simple dynamic walking model predicts the mechanical work performed on the body center of mass (COM) to redirect the COM velocity between steps. (left to right in B:) Level flat walking is powered by active push-off (PO) work from the trailing leg to offset collision losses (CO) from the leading leg, to redirect velocity *v*^−^ from the end of one arc to *v*^+^ at beginning of next. Upward and downward steps require different amounts of work to maintain same speed, where additional work may also be performed about the stance hip (in gray) to gain speed. (C) Model predicts work rates should increase with walking speed (*v*^3^) and terrain amplitude changes (*h*^2^). Metabolic rate is predicted to increase in approximate proportion to the work rate, due to active work by muscle.

Interestingly, these various control strategies predict the same, testable dependence on walking speed and terrain amplitude [22]. Whether with optimal anticipatory planning or constant push-off or hip work, the positive work *W*_com_ on the COM per step increases with speed *v* and terrain amplitude *h* approximately with the power law

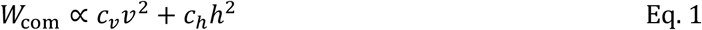

where *c*_*v*_ and *c*_*h*_ are empirical coefficients and *h* is the maximum amplitude of random step heights, measured peak-to-peak. The coefficients can differ markedly depending on the control strategy, which we do not predict, but the power law relationship is a testable prediction for the human’s COM work as a function of speed and terrain. The speed dependence is because the model’s positive work per step is determined by the impact speed and inter-leg angle, increasing in approximate proportion to the square of walking speed *v* [19]. The terrain dependence is because similar COM redirection mechanics are important even on uneven terrain. The quadratic term may be explained by the fact that *W*_com_ should be an even function of *h*, because random terrain is statistically invariant to the sign of *h*, so that the square power must be the leading term of a polynomial expansion for *W*_com_.

Energy expenditure is hypothesized to increase in rough proportion to the work rate. The work performed by muscle has long been observed to contribute approximately linearly to, and account for most of, overall energy consumption [23], [24]. A very simple prediction is that COM work and energy rates increase approximately proportionately [13]:

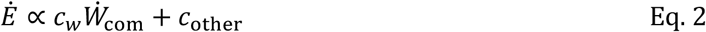

where coefficient *c*_*w*_ is inverse efficiency, *W*_com_ is the positive work rate (work per stride for one leg multiplied by step frequency), and *c*_other_ is for costs independent of work. A typical benchmark efficiency (work rate divided by metabolic rate) is about 25% [12], but overall efficiency may be lower if energy is expended for physiological processes not due to work [23], and higher if series elastic tissues perform substantial work passively, as has been proposed for level walking and running [25]. Our prediction is concerned only with increases in work and energy, quantified by *c*_*w*_ or its inverse, a delta efficiency. We have no specific expectation for the delta efficiency, due to likely elastic contributions [25], and therefore test mainly for the linear proportionality. The final term *c*_other_ lumps together a number of other physiological costs not modeled here, such as for moving the legs back and forth with respect to the COM [26], [27], maintaining balance, and producing force without work.

Another prediction is that energy cost should increase with walking speed and terrain amplitude. This follows from combining the work and proportionally predictions (Eqs. 1 and 2) and adding a simplification, that speed increases in proportion to step frequency, resulting in the prediction:

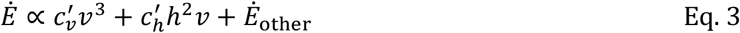

Human speed also increases with step length as well, but less so than with step frequency [28] and predicting a slightly higher exponent with *v*^3.42^[4], a difference too small to be resolvable here.

### Human subjects experiment

We measured healthy young adults walking on a split-belt instrumented treadmill (Bertec, Columbus, Ohio, USA) modified for uneven terrain. Ten subjects participated (N = 10, six females and four males, age 27.69 ± 5.21 yrs). Energetics data were collected for four terrain amplitudes *h* (0, 0.019, 0.032, 0.045 m), each at four speeds *v* (0.8, 1.0, 1.2, 1.4 m · s^−1^), for a total of sixteen unique conditions. Amplitude *h* is defined as a peak-to-peak maximum amplitude of uneven heights of the walking surface. The terrains were applied in random order, and the speeds for each terrain in random order. Before the experiment, subjects provided written informed consent, as approved by the University of Calgary Board of Ethics.

The uneven terrain treadmill has been described previously ([18], inspired by [1]) and is only briefly summarized here (Figure 2A). A standard treadmill belt was affixed with synthetic uneven terrain, consisting of strips of polystyrene construction insulation that could pass over the treadmill rollers unobstructed. The strips were arranged in blocks 0.3 m long in the walking direction, with a random but set of discrete amplitudes within the limit *h* for each terrain condition (Figure 2A). There were two such treadmill belts, offset by a random fore-aft distance to reduce predictability of step heights. Each footfall typically occurred on either a flat block or across an upward or downward transition between blocks. We also applied an additional “Rough” terrain of more continuous small irregularities, not of polystyrene but rather dried construction glue of approximately 0.005 m amplitude. Because this surface was qualitatively different from the others, the data are included in some figures for illustration only but were not used in any quantitative analysis across discrete amplitudes.

**Figure 2:**
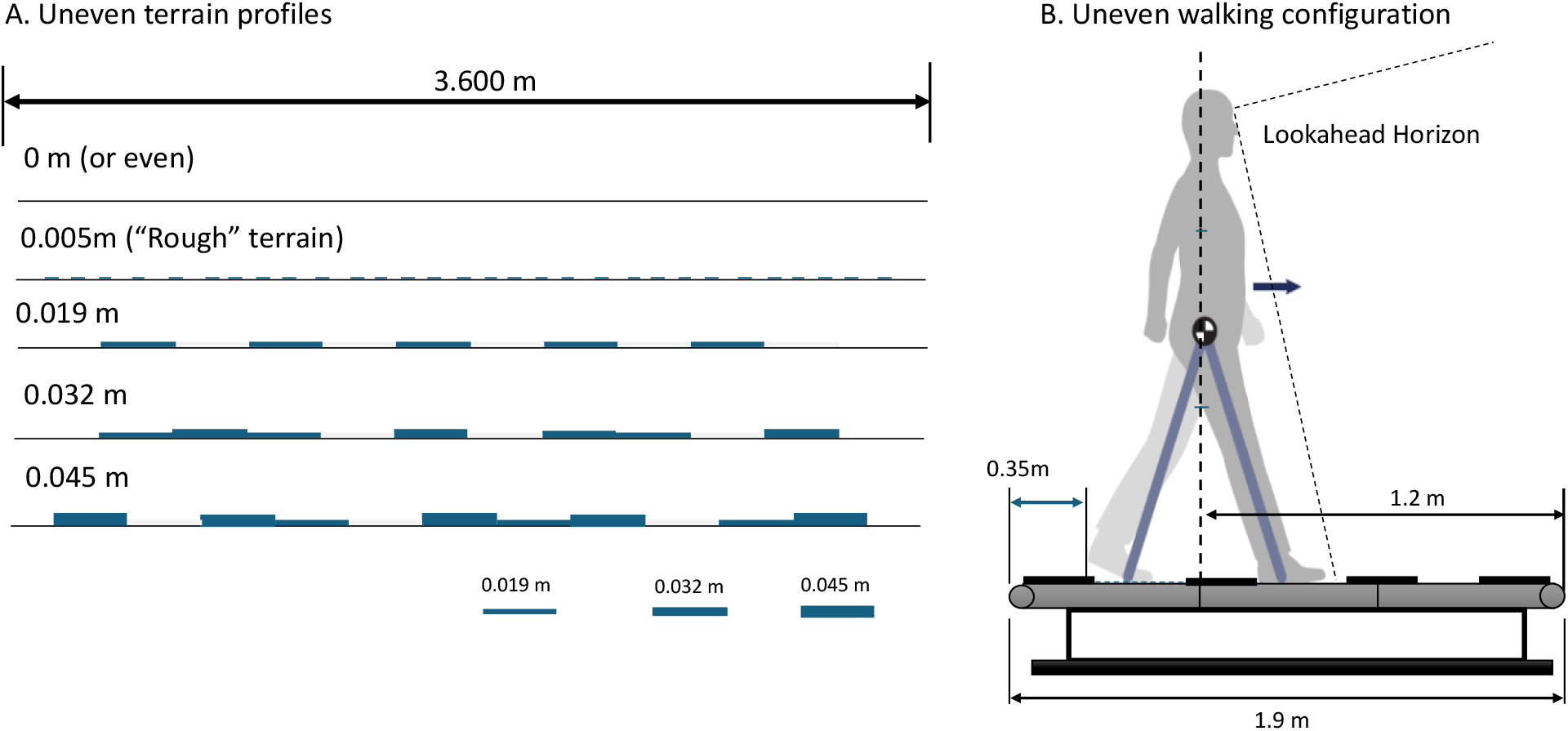
Experiment with uneven walking terrain. (A) Uneven walking terrain profiles, with peak-to-peak amplitudes of 0 to 0.045 m. Terrain was constructed from strips of polystyrene arranged into blocks 0.3 m long and as wide as one treadmill belt, with two such belts on a split-belt instrumented treadmill. (B) The walking configuration placed the subject with about 1.2 m (nearly two step lengths) of lookahead horizon to see the coming terrain. Fore-aft position was cued by a slack rubber tubing placed across the treadmill and gently contacting the subject’s waist.

Data collection included ground reaction forces and respirometry, along with limited motion capture. Ground reaction forces and moments for each leg were collected at 960 Hz (low-pass filtered, 10 Hz). These forces were utilized to detect step times [29] and were also integrated to estimate body center of mass (COM) velocity [13] with a high-pass filter at an ad hoc 0.26Hz cut-off to reduce drift. Oxygen consumption and carbon dioxide production rates were recorded using the Cosmed K5 metabolic system (Cosmed S.r.l., Rome, Italy). Trials were six minutes long, with initial transients discarded and only the last three minutes used in standard equations [30] to determine steady-state net metabolic rates. We subtracted each subject’s average standing metabolic rate (1.44 ± 0.23 W · kg^−1^ collected separately before walking trials) from the gross rates. Whereas forces and respirometry data were collected at all sixteen combinations of the four speeds and four terrains, motion capture (Phase Space Inc, CA, USA, sampling rate 960 Hz) was also collected for only seven combinations: all terrains at one nominal speed (1.2 m · s^−1^), and all speeds at one nominal amplitude (0.032 m). A standard marker set [1] was collected, of which only a small subset was used here to determine gait parameters, from foot and the pelvis markers. Each foot was measured with a heel (calcaneus) marker, and a metatarsal phalangeal joint (MTP) from the spatial average of first and fifth metatarsal markers.

Subjects could see the upcoming terrain up to two steps of ahead. We attempted to regulate a home position about 1.2 meters behind the front of the treadmill belt (Figure 2B), marked by slack surgical tubing hung laterally across the treadmill. We instructed subjects to look ahead actively and to approximately maintain home position by feel, from gentle contact between torso and tubing. They were occasionally reminded through verbal feedback if they deviated substantially from home. Before the walking trials, subjects practiced for a few minutes to become familiar with the treadmill and home position.

We assessed kinematic gait parameters such as step length and width, and kinetic measures such as COM work, as well as respective variabilities. Step length and width were estimated from heel marker positions projected on the treadmill belt at consecutive heel strikes. A virtual foot clearance was estimated from the distance between a local minimum in MTP height mid-swing and an assumed terrain height linearly interpolated from toe-off to the subsequent heel-strike instances [2]. Variabilities were quantified as root-mean-square (RMS) variability across the discrete steps for one minute of data for each condition.

Kinetic measures were derived from instantaneous COM power, the dot product of COM velocity and ground reaction force [31]. The COM velocity was determined by integrating ground reaction forces, with the aforementioned high-pass filter to reduce integration drift. Positive and negative work per stride were determined by integrating the positive and negative power intervals. Similarly, push-off and collision work per stride were integrated from the respective intervals about double support [31].

We tested model predictions for work and energy expenditure. The primary mechanistic predictions were that positive work per step would increase with speed and terrain (Eq. 1), and that net metabolic rate would increase with speed and terrain (Eq. 3), tested with linear regression for dependencies on *v* and *h* raised to appropriate exponents. We defined a nominal uneven terrain (*h*=0.032 m) for summarizing effects of increasing speed, and a nominal speed (1.2 m · s^−1^) for amplitude effects. We tested for a linear proportionality irrespective of speed and terrain, between differences in energy rate and work rate (*c*_*w*_ in Eq. 2), or its inverse (delta efficiency 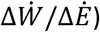. Positive work rate was calculated by dividing the positive COM per step by the associated step duration [32]. These regressions included individualized offsets for each subject’s work or energy across all conditions, implemented with linear mixed effects models (statsmodels 0.15.0 [33]) with subject treated as a random categorical effect. These offsets were included in all statistical metrics but were subtracted from energetics plots to deemphasize intersubject variability.

We also tested for statistical relationships for kinematic and other variables. Kinematic gait parameters and work variabilities were tested for linear dependencies on speed and amplitude conditions, using ordinary least squares. All tests were reported with regression coefficients (gains), *R*^2^ (including conditional *R*^2^ for linear mixed models [34]), and P-values with significance threshold 0.05.

All analyses were performed on non-dimensionalized data, using subject mass *M* (71.00 ± 6.16 kg), leg Length *L* (0.926 ± 0.033 m, floor to greater trochanter), and gravitational acceleration *g* as base units. For work or energy, the average normalization factor *MgL* was 644.97 ± 80.94 J, and for energy rates *Mg*^1.5^*L*^0.5^ was 2099.26 ± 222.43 W. Discrete speed conditions were non-dimensionalized by average *g*^0.5^*L*^0.5^, and terrain amplitude by average *L*. Dimensionless data were used to facilitate averaging and comparison across subjects of different sizes and reported in tabular results. All plots are of dimensionless data but labeled with more traditional dimensional axes using the average normalization factor.

## Results

Walking on uneven terrain caused multiple changes as a function of walking speed and terrain amplitude. These included qualitative effects, such as relative smooth changes to ground reaction forces. For example, the vertical GRF (Figure 3) and COM power profiles resembled those for flat walking but had peaks that appeared to grow with walking speed. Greater terrain amplitude appeared to alter the trajectories further, for instance, with greater initial force peak and lesser second peak. The COM powers appeared to increase in amplitude, and perhaps in timing with terrain changes (Figure 4). These informal observations are followed by more quantitative results below.

**Figure 3:**
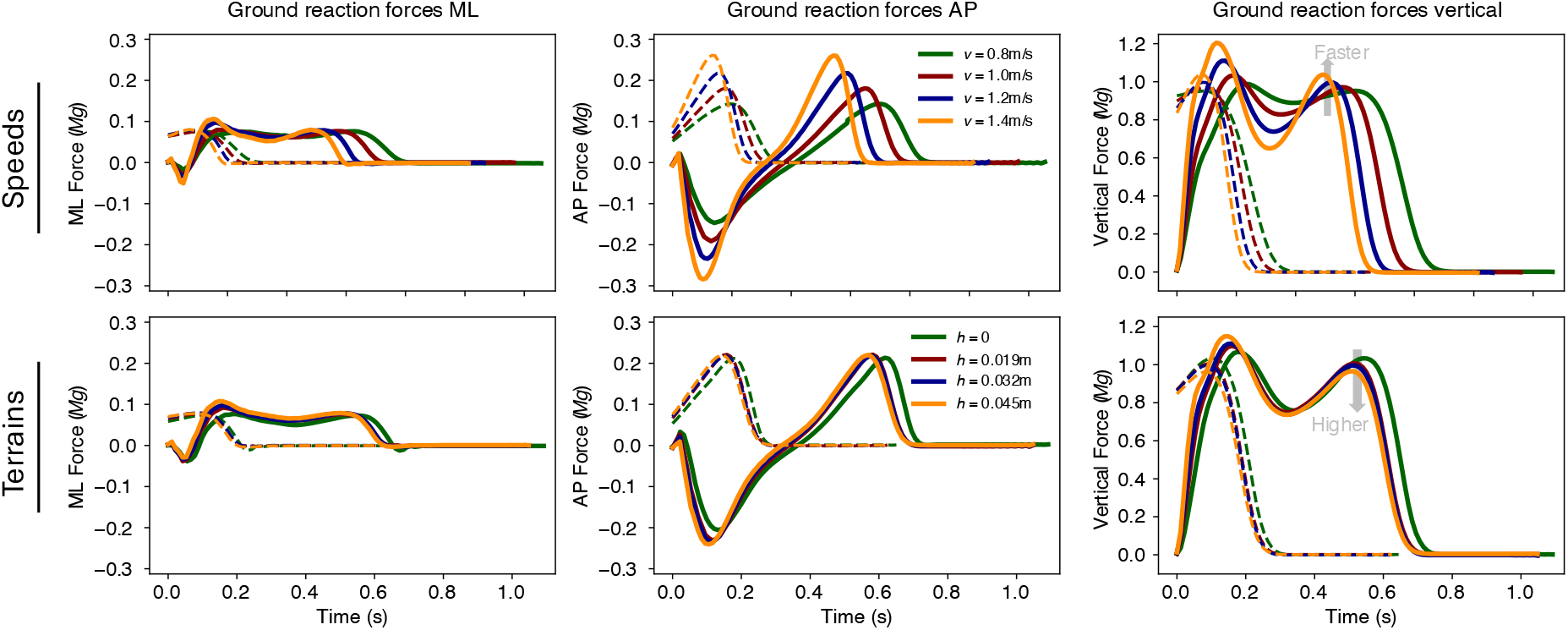
Ground reaction forces vs. time for (top row) varying walking speed and (bottom row) varying terrain amplitude. Speeds *v* are 0.8, 1.0, 1.2, and 1.4 m · s^−1^, and amplitudes *h* are 0, 0.019, 0.032, 0.045 m peak-to-peak. Forces are shown averaged across about 50 strides from each leg and across subjects (solid lines); other leg is also shown during double support (dashed lines).

**Figure 4:**
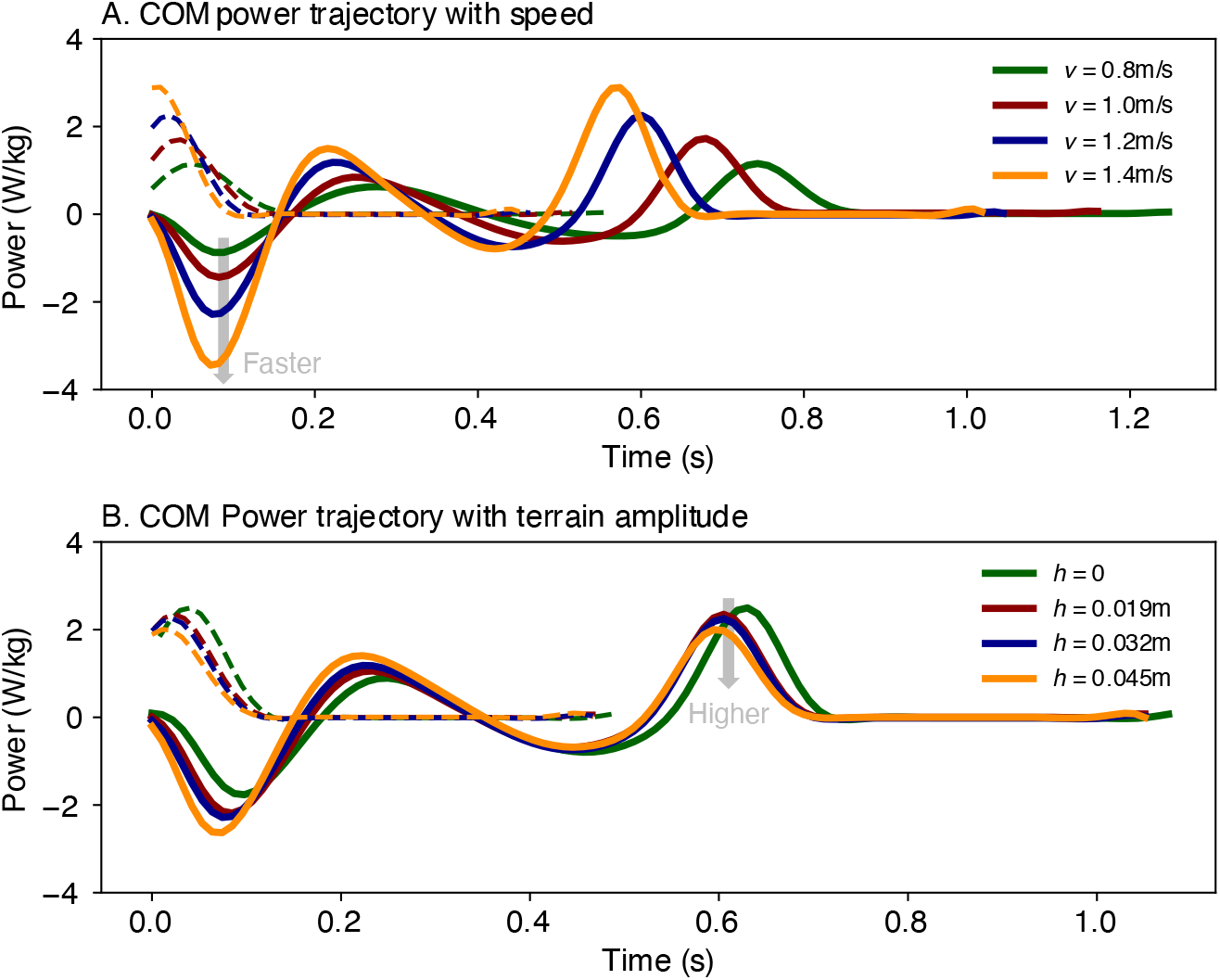
Mechanics of walking on uneven terrain, in terms of COM work rate vs. time for varying (A) walking speed *v* (0.8, 1.0, 1.2, and 1.4 m · s^−1^) and varying (B) terrain amplitude *h* (0, 0.019, 0.032, 0.045 m) peak-to-peak. COM powers are the dot product of individual leg ground reaction force and COM velocity, averaged across about 50 strides per subject and across subjects (solid lines); other leg is also shown during double support (dashed lines).

Walking speed and uneven terrain affected kinematic gait parameters in several respects, determined statistically by linear relationships (Figure 5). Step length and virtual clearance both increased with walking speed, stride duration decreased, and step length and duration (root-mean-square, RMS) variabilities changed with speed (Table 1). Focusing on terrain amplitude, step width and virtual clearance both increased, by 3.5%/cm and 18%/cm, respectively (in terms of linear sensitivities per 1 cm of terrain amplitude). All of the corresponding variabilities increased, by 22%/cm for step length, 14%/cm for step width, 76%/cm for virtual clearance, and 25%/cm for step duration. Thus, the most notable effects of uneven terrain were on virtual clearance and its variability, as well as gait variability in general.

**Table 1:**
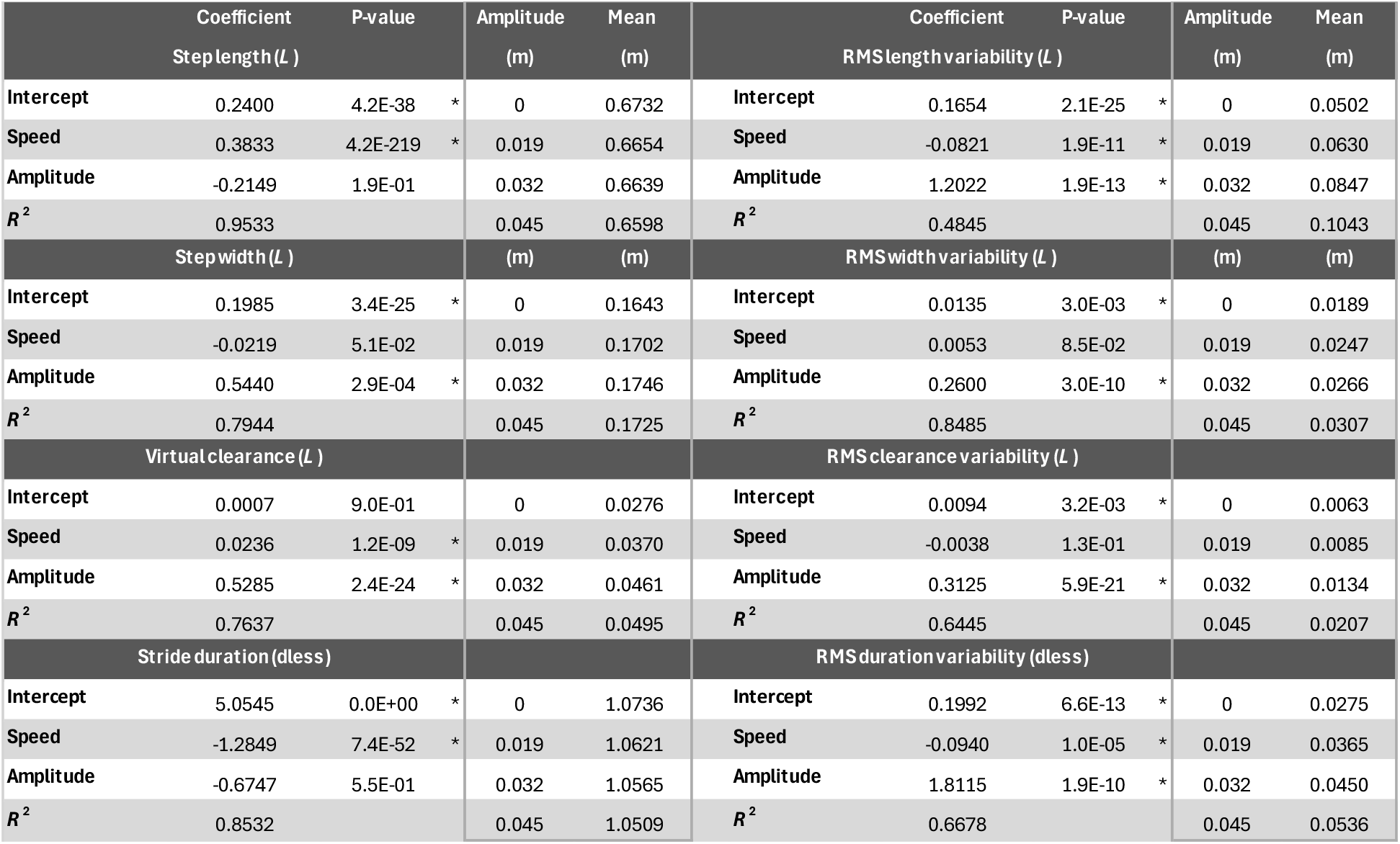
Sensitivities of kinematic gait parameters to walking speed and terrain amplitude. The dependent variables were step length, step width, virtual foot clearance, and stride duration (left column), along with root-mean-square (RMS) variabilities. Each was tested as a statistical linear model, yielding an intercept and coefficients for speed (m · s^−1^; at nominal amplitude 0.032 m) and terrain amplitude (m; at nominal speed 1.2 m · s^−1^), with asterisk (*) indicating significance P < 0.05. For reference, the mean values for each terrain amplitude are shown (boxed area; at nominal speed 1.2 m · s^−1^). Kinematic data were collected for four terrain amplitudes at nominal speed and four speeds at nominal amplitude, for a total of seven unique conditions.

**Figure 5:**
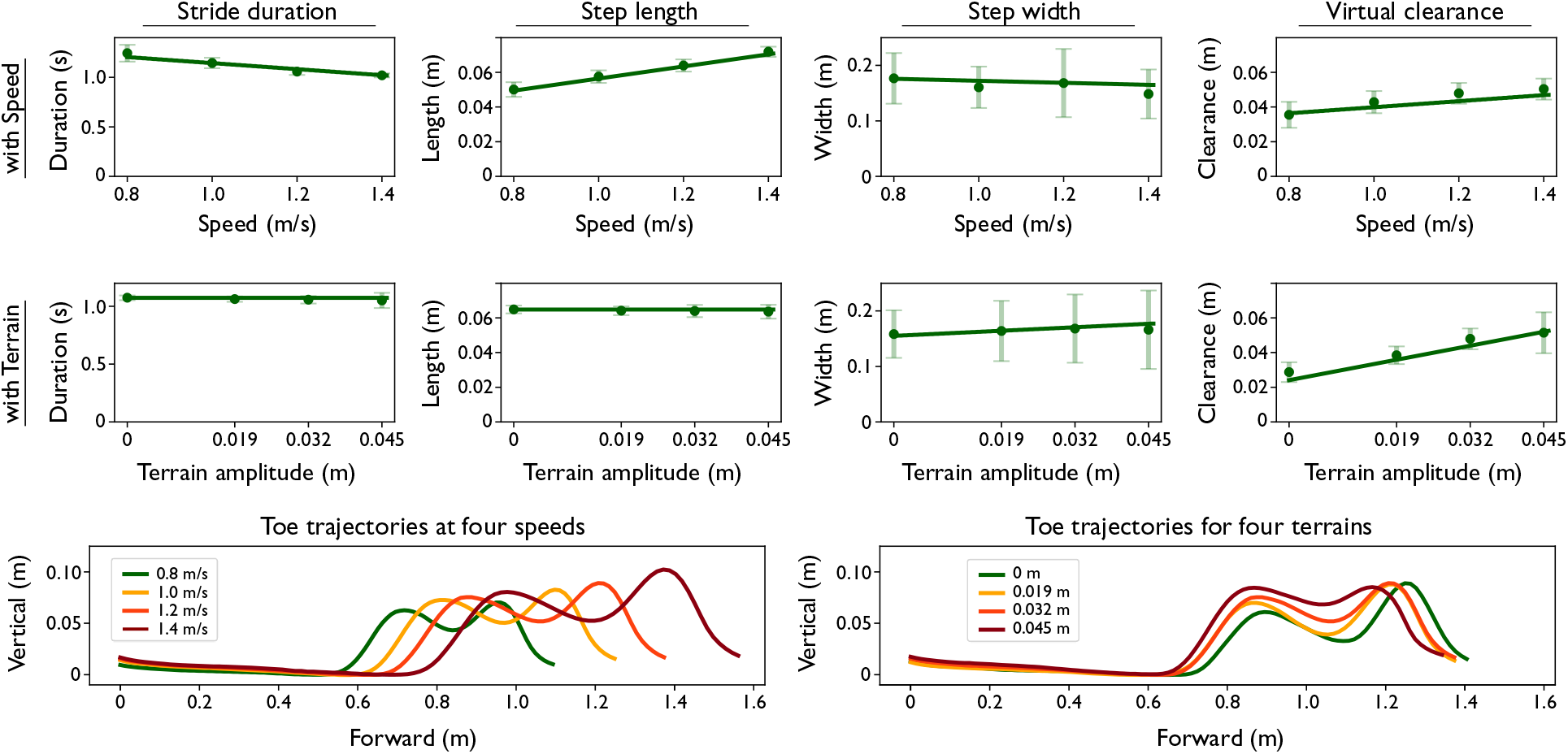
Kinematic gait parameters as function of walking speed and terrain amplitude. The quantities shown (mean ± s.d., filled symbol and error bars) are (left to right columns:) stride duration, step length, step width, and virtual clearance, as functions of (first row:) walking speed at fixed terrain amplitude 0.032 m, (second row:) and terrain amplitude at fixed speed 1.2 m · s^−1^. (third row:) Average trajectories for toe motion in the sagittal plane (vertical vs. forward displacement, relative to heel strike).

For kinetics, the positive and negative COM work magnitudes increased with speed and terrain amplitude squared (Figure 6). The overall fits for work agreed reasonably well with predictions (Eq. 2), with *R*^2^ ranging 0.67 − 0.78 (Table 2A). Based on fits, the positive and negative work magnitudes per step approximately doubled with speed, increasing by 95% and 100%, respectively over the speeds examined on nominal terrain (*h* = 0.032 m). The work magnitudes also increased with terrain amplitude, albeit less substantially, by 17% and 14%, respectively over the amplitudes examined at nominal speed (1.2 m · s^−1^). Push-off work per step increased by about 153% across the speeds examined, and collision work magnitude by about 282%. With increasing terrain amplitude, push-off per step increased negligibly by 2.4%, whereas collision work magnitude increased more substantially, by about 27%.

**Table 2:**
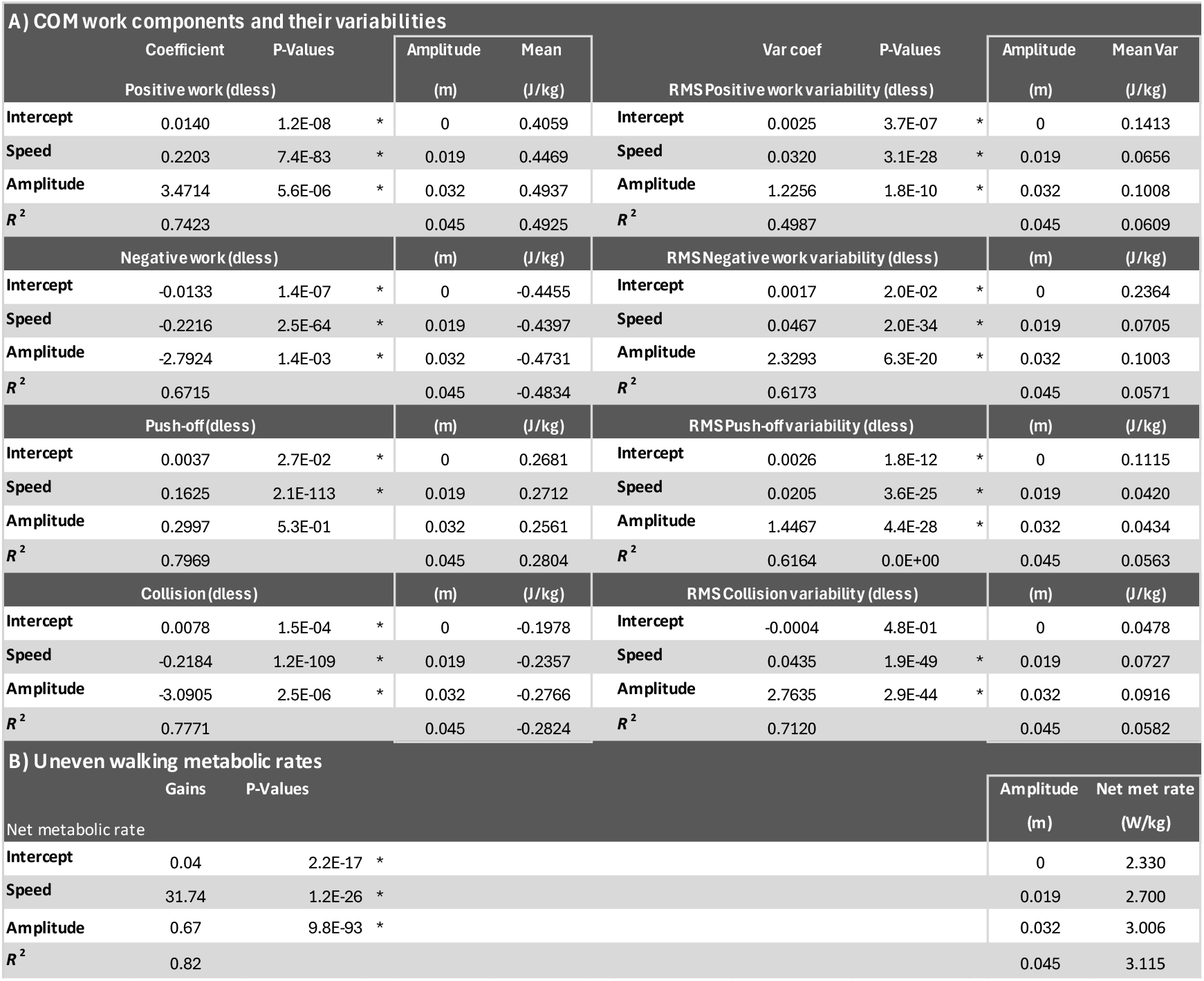
Sensitivities of COM work and net metabolic rate to walking speed and terrain amplitude. The dependent variables were (left column) total positive work, push-off work, total negative work, and collision work (per step for all), along with their respective RMS variabilities (right column). The work quantities were tested against model predictions of speed squared and terrain amplitude squared (Eq. 1), and RMS variabilities against linear speed, using statistical linear models. For energetics, the dependent variable was net metabolic rate, and the independent variables were speed cubed and terrain squared (Eq. 3). These yielded intercept and coefficients, with asterisk (*) indicating significance P < 0.05. For reference, the mean values for each terrain amplitude and nominal speed 1.2 m · s^−1^ are also shown (boxed area).

**Figure 6:**
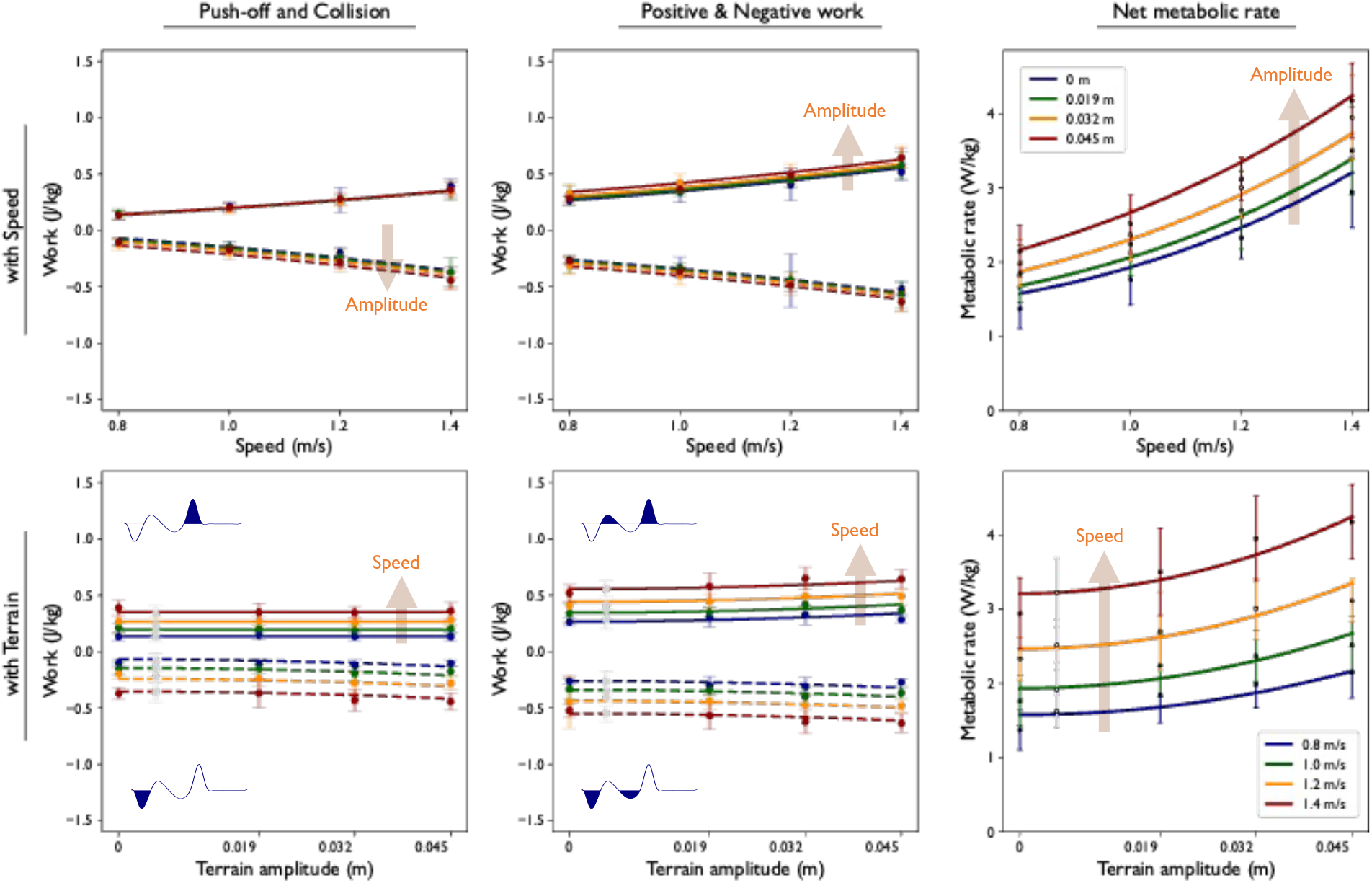
Work and energy costs for walking with (top row:) increasing speed and (bottom row:) increasing terrain amplitude. Work is shown as (left to right:) push-off and collision per step, total positive and negative work per step, and energy as net metabolic rate. Experimental data (N = 10, mean ± s.d. error bars) are shown for each condition, along fitted curves according to model predictions: Work magnitudes increasing with speed squared and amplitude squared (*W* = *c*_*v*_*v*^2^ + *c*_*h*_*h*^2^ + *d*), and metabolic rate with speed cubed and amplitude squared 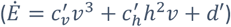.Positive and negative work curves are distinguished by solid and dashed lines, respectively. Push-off and collision are defined as the respective positive and negative work intervals encompassing double support, and positive and negative work for the entire step for one leg (illustrated by inset figures). All of the coefficients were statistically significant (P < 0.05), except for push-off with terrain amplitude; see Table 2 for details.

This suggests that positive work was performed differently with increasing speed than with increasing amplitude. Similar to previous reports [13], subjects walked faster mostly by increasing push-off and collision. Specifically, push-off accounted for about 74% of the positive work increases with speed, and collision for 99% of the negative work changes. But with increasing terrain amplitude, push-off accounted for only 8.6% of the positive work changes, whereas collision accounted for 111% of the negative work changes. In other words, the greater positive work on uneven terrain was almost entirely *not* attributable to push-off, but rather 91.4% attributable to what we call rebound ([31], [35]) following collision. The greater work for walking on uneven terrain was largely performed not at push-off, which is mostly about the ankle [31], but rather to some combination of stance hip and knee [1].

Work variabilities increased substantially and approximately linearly with walking speed and terrain amplitude (Table 2A). From statistical fits, the total RMS positive and negative work variabilities increased by about 74% and 86% respectively with speed (at nominal amplitude), and 44% and 70% respectively with amplitude (at nominal speed). Similarly, push-off and collision work variabilities increased by about 50% and 102% with speed, and 67% and 120% with amplitude.

The net metabolic rate also increased with walking speed and terrain amplitude (Figure 6). The cost agreed reasonably well with model predictions of speed cubed and terrain amplitude squared, with *R*^2^ of 0.82 (Table 2B). The fits were equivalent to a near doubling of metabolic rate (99% increase) with speed (at nominal amplitude 0.032 m), and a 36% increase with amplitude (at nominal speed 1.2 m · s^−1^. Expressed in dimensional units, the metabolic rate coefficients were equivalent to 0.732 W · kg^−1^ · s^3^ · m^−3^ for speed and 366 W · kg^−1^ · s · m^−3^ for amplitude.

Both positive work and net metabolic rates increased in proportion with each other across conditions (Figure 7). Each agreed reasonably well with the same form of model prediction (Eq. 3), shown as a surface function of walking speed and terrain amplitude (Figure 7A and B for work and metabolic rates, respectively). This suggests that the changes should also be proportionate. Indeed, metabolic rate increased approximately in proportion with positive work rate (Figure 7C), yielding a slope of 2.019 (95% CI: 1.735, 2.302; P = 2.7e-44, *R*^2^ = 0.67; linear mixed model with individual intercepts treating subjects as categorical random effect). This was equivalent to a delta efficiency 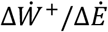 of 49.5%. From their approximately linear relationship, work could thus explain about 67% of the variance in metabolic rate across conditions. There was also a significantly non-zero intercept in the energy-work relationship, 1.01 W · kg^−1^ (*P*=1.4e-8) or about 34% of the net metabolic rate for nominal uneven walking.

**Figure 7:**
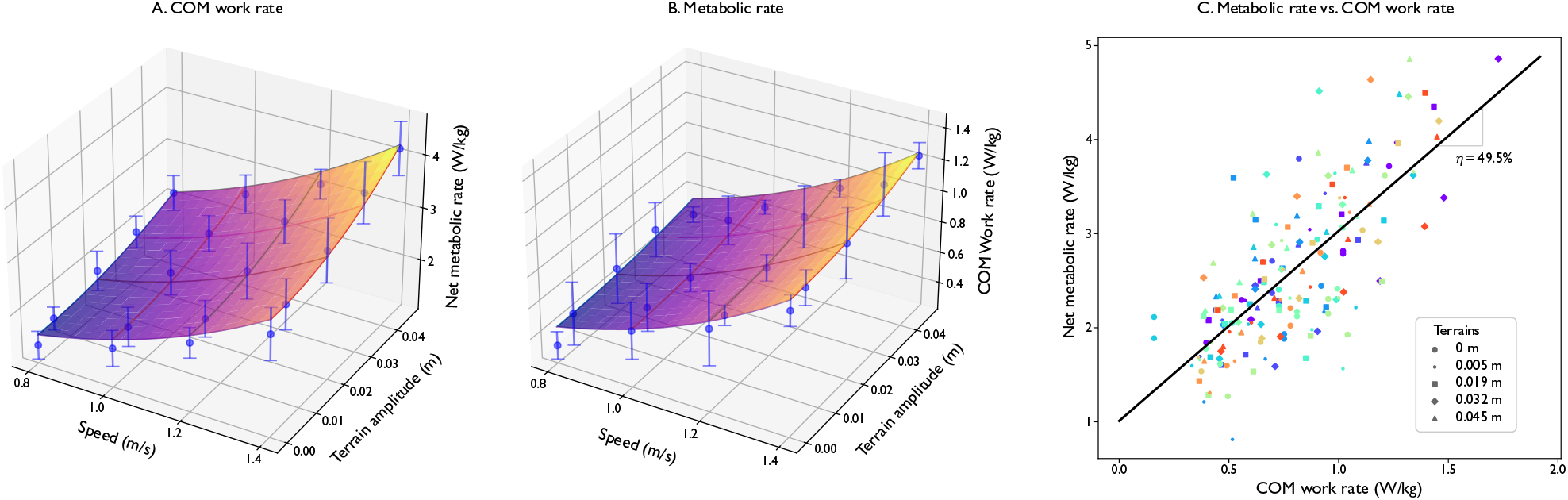
Work and net metabolic rate plotted for all conditions (N = 10), and against each other. (A.) Positive COM work rate and (B.) net metabolic rate as a function of four walking speeds and four terrain amplitudes, along with corresponding model fits (Eq. 3). (C.) Net metabolic rate vs. positive COM work rate for all conditions, along with a linear fit yielding a delta efficiency (inverse slope) η = 49.5%. Plots also include rough terrain condition at approximate amplitude 0.005 m, not included in model fits. Data and fits are the same as Figure 6 and Table 2, except (A.) work rate (*R*^2^ = 0.74) is plotted rather than work per step, yielding units comparable to metabolic rate (*R*^2^ = 0.82). Filled symbols denote means across subjects, error bars denote standard deviation after removing individualized offsets. The fit in (C.) is linear mixed model, yielding slope 2.065 (P=2.7e-44) and intercept 1.01 W/kg (P=1.4e-8), *R*^2^ = 0.67.

## Discussion

We had sought to parametrically characterize the effect of uneven terrain on metabolic energy expenditure. This included the effect of terrain amplitude on work performed on the COM, gait parameters, and respective variabilities. We found that metabolic rate was explained reasonably well by the simple kinetic measure of COM work, in agreement with a simple mechanistic model. This also made it possible to predict energetics as a function of condition, namely walking speed and terrain amplitude. These results suggest that the energetics uneven walking have similar determinants as flat walking, with costs varying parametrically, rather than discretely, with terrain.

The mechanistic model predicted how both speed and terrain affect mechanical work production. The observed increase in COM work per step, approximately with speed squared, replicates previous tests for flat walking [31], [36], [37], explained by redirection of COM velocity. But previously untested was the effect of terrain, hypothesized to be amplitude squared (Eq. 1), because uneven steps require more redirection work. Both speed and terrain-dependent observations agreed well with data (Figure 6A and B) according to the mechanistic model (Eq. 1). Whereas previous studies have generally treated uneven terrain as a discrete condition, we found that terrain amplitude may be treated as a continuous parameter increasing smoothly from zero for flat walking. This provides a consistent mechanistic basis for understanding how humans walk faster or on uneven terrain.

This work had an approximately proportional effect on energy expenditure. Metabolic rate increased approximately with speed cubed and amplitude squared (Figure 6), as expected if muscles expend energy proportional to active work (Eq. 2). Indeed, a direct comparison revealed an approximately linear relationship between metabolic and COM work rates (Figure 7), where changes in work could explain about 67% (from *R*^2^) of the variance in metabolic rate across all conditions. We did not have a specific prediction for the associated coefficient, which was equivalent to a delta efficiency of 49.5%. This is roughly within the range of previous delta efficiencies estimated for flat walking and running [25]. Delta efficiencies that exceed the 25% typical for positive muscle work [12] have traditionally been attributed to passive work by series elastic tissues [25]. Here we evaluate work for the two legs separately [38] rather than combined [25], because the energy cost of positive work of one leg is not plausibly reduced by the negative work of the other. We nonetheless agree with the interpretation of elastic contributions, having speculated about elastic rebound following collision [31] and modeled elastic push-off following preload [39]. A single delta efficiency seems applicable across conditions, suggesting that energy cost may have similar work dependence irrespective of speed or terrain.

The mechanical work was, however, performed differently with increasingly uneven terrain. Positive work increased almost entirely during early single stance (rebound phase) with terrain amplitude, as opposed to later (during push-off) with faster walking (Figure 6). We had no specific prediction for the amplitude-squared coefficient for work (Eq. 1, *c*_*h*_), which could theoretically be quite small with perfectly pre-emptive push-off timing before collision [15], because the ideal push-off reduces collision losses [19] and is at least partially elastic [39]. But the coefficient could also be quite large if work is not performed at push-off or if the push-off is late or not anticipatory [16], [22], resulting in greater collisions. Humans do appear to anticipate well when walking overground [21], but perhaps less so on a treadmill, where there is restricted view of upcoming terrain and an artificial speed constraint due to the need for station-keeping [40], [41]. Some of the greater work observed here may therefore be attributable to treadmill conditions. However, overground walking data also show greater work, primarily at stance hip and knee and associated with a more crouched posture [1], agreeing with the present rebound observations. We also suspect that uneven terrain contributes to poorer timing and elastic return for push-off. Work performed on the COM does explain much of the high cost of uneven walking, but there is more to be discovered about how that work is performed.

There was also significant energy expenditure *not* explained by changes in COM work. Net metabolic rate (Figure 7) with respect to work (Eq. 2) had a non-zero intercept, equivalent to about 34% of the energy expenditure for nominal uneven walking, as well as unexplained variance of about 33%. Net metabolic rate curves usually have non-zero intercept with respect to speed [3], and our results show a similar intercept for speed and amplitude combined. The unexplained variance we observed is attributable in part to within-subject variability, and perhaps to other deterministic effects not captured here.

Other measures may be indicative of such unmodeled energetic costs. Of the kinematic gait parameters and variabilities examined, we found virtual clearance and all gait variabilities to increase substantially with terrain (e.g., at least 10%/cm). There were increases in work variability (Table 2), suggesting that unsteady steps are associated with unsteady muscle force and work, perhaps leading to greater energy cost [42]. Indeed, measures such as virtual clearance and gait variability have also been observed to increase with (qualitative) unevenness of natural terrain and associated energy cost [2]. Controlled experiments have also shown that greater virtual clearance drives greater energy expenditure [43], as does greater gait variability induced by visual perturbations [6], [7]. Some of these effects could potentially be quantified by kinetic measures not examined here, such as joint torque or power, or work peripheral to the COM [37]. This could potentially provide a mechanistic basis for deterministic energy costs not related to COM work, such as for lifting the swing leg, adjusting foot placement, or otherwise stabilizing balance. Such costs would ideally be evident in unexplained biases in the energy vs. COM work relationship, but are unfortunately not obvious in present data (Figure 7).

This study advances the ability to examine uneven walking parametrically and mechanistically. Previous studies have treated terrain discretely, for example as a terrain coefficient [17] or experimental condition [1], [2], [5], but with little ability to either interpolate between conditions or to understand them mechanistically. We introduced terrain amplitude as a continuous parameter that could be incorporated in a mechanistic model and a parametric experiment. The uneven walking model has the same pendulum-like and step-to-step mechanics as previous models of steady flat walking [19], except with varying speed and step height [15]. The same model can therefore walk on flat ground as a special case of uneven walking, and at other speeds. Experimentally, terrain amplitude can be known in advance along with speed and used predictively. And with a split-belt instrumented treadmill, it was possible to test both mechanics and energetics predictions from the model.

The primary limitation of the present study was a relatively narrow range of terrains. The inverted pendulum model only applies to a small range of terrain amplitudes where collisions dissipate significant energy and yet there is substantial momentum retained between footfalls. For larger amplitudes, beyond say the height of a sidewalk curb, it is sensible to land with a flexed knee with less dissipative collision, and more of the positive work performed by that same leg than by trailing leg push-off. The experiment was similarly limited to small terrain amplitudes [18], and so neither model nor data inform what happens at greater terrain amplitude. Small amplitudes may also explain why we detected only linear effects between speed and terrain, in contrast to the terrain coefficient, which is multiplicative with speed [17]. But real-world terrain also varies in many ways beyond amplitude alone, for example softness, compactness, or squishiness. Our results are thus confined to firm terrain with only modest unevenness.

There were several additional limitations. We only examined relatively simple measures of gait and energetics, as averages or gait variabilities. If humans perform active feedback corrections [44], [45], the mechanics and energetics would be better quantified as causal, state-dependent actions both between and within steps [46]. Such actions also presumably depend on sensory feedback, some of it measurable such as visual gaze [35]. We also did not perform detailed examination of joint kinematics or kinetics. Analyses such as inverse dynamics might quantify forces and work other than on the COM, such as for foot placement. We also limited the present analysis to mechanistic relationships informed by model, whereas a more intensive statistical (and less mechanistic) approach [2] could potentially identify other explanatory relationships.

Humans perform more work and expend more energy as they walk faster or on more uneven terrain. The energy rate increases approximately with the cube of walking speed and the square of terrain amplitude, approximately in proportion to the rate of work performed on the COM. A simple model predicts the work needed to redirect the COM velocity between pendulum-like steps, which can proportionally explain about two-thirds of the metabolic cost. We also suspect that work is performed on the COM differently on uneven terrain, and energy expended for other actions such as to adjust foot placement or correct balance contribute to the higher cost for uneven terrain. Although such costs have yet to be quantified, but much of the overall energetic cost of walking faster or on uneven terrain is mechanistically explained by greater active work on the COM.

## Data and resource availability

The dataset for this study will be made available in a publicly accessible archive.

